# Inhibition of p107 alleviates liver steatosis by reducing de novo fatty acid synthesis

**DOI:** 10.64898/2026.04.14.718271

**Authors:** Juan Cunarro, Marta V. Miguens, Tadeu de Oliveira-Diz Tadeu, Xabier Buque, Lucia Oro, Cristina Riobello, Jose Iglesias, Carmen Quintela-Vilariño, Ana T. Maduro, Alba Cabaleiro, Eva Novoa, Alejandro Fuentes-Iglesias, Miguel Fidalgo, Diana Guallar, Anxo Vidal, Alfonso Mora, Marta Varela-Rey, Ruben Nogueiras, Guadalupe Sabio, Patricia Aspichueta, Lluis Fajas, Carlos Diéguez, Sulay Tovar

**Author notes:** **Corresponding author:** Dr. Sulay Tovar, *Diabesity group* CIMUS (Centro de Investigación de Medicina Molecular y Enfermedades Crónicas) Universidade de Santiago de Compostela, Avda Barcelona s/n, 15782 Santiago de Compostela, Spain. Contributed equally to this work (Juan Cunarro, Marta V. Miguens).

## Abstract

Metabolic dysfunction–associated steatotic liver disease (MASLD) is characterized by excessive hepatic lipid accumulation driven by increased de novo lipogenesis (DNL) and impaired lipid oxidation. p107, a member of the retinoblastoma (Rb) family, extensively studied in the context of cell cycle regulation and adipocyte differentiation recently has been identified as a metabolic regulator controlling thermogenic activity. However, its role in hepatic lipid homeostasis remains poorly understood. Here, we identify the cell cycle regulator p107 as a key modulator of hepatic lipid metabolism.

p107 expression is increased in patients with MASLD and correlates with disease severity. In mouse models, global and liver-specific p107 deficiency protect against high-fat diet-induced steatosis without affecting body weight. This is associated with reduced expression of lipogenic enzymes including fatty acid synthase (FASN), and enhanced mitochondrial oxidative pathways. Conversely, hepatic restoration of p107 reversed these effects and promoted lipid accumulation and endoplasmic reticulum stress. Consistent with this in human hepatocytes, p107 silencing reduces lipid accumulation, decreases DNL and enhances mitochondrial respiration, whereas p107 overexpression induces the opposite phenotype. Notably, FASN knockdown attenuates the pro-steatotic effects of p107, indicating that it is a critical downstream mediator of p107.

Together, these findings establish p107 as a physiological regulator of hepatic lipid metabolism, with its dysregulation contributing to the development of MASLD.

## 1. Introduction

Metabolic dysfunction-associated fatty liver disease (MASLD) is the most common cause of chronic liver disease in the Western world and is rapidly increasing in prevalence globally. MASLD is a frequent comorbidity of type 2 diabetes and obesity, with its prevalence calculated at ∼30% in the general population and 80% among obese people. MASLD encompasses a disease spectrum ranging from simple steatosis, characterized by triglyceride accumulation in hepatocytes to metabolic dysfunction-associated steatohepatitis (MASH). The hallmarks are inflammation and fibrogenesis, which can further progress into cirrhosis and hepato-cellular carcinoma (HCC), the deadliest form of liver cancer (Tilg and Moschen, 2010). In the other hand although it was generally assumed that only MASH progressed to cirrhosis and end-stage liver disease, recent studies indicate that MASLD can also lead to progressive fibrosis (Pais et al., 2013; Schattenberg and Schuppan, 2011; Ye et al., 2020).

The progression of MASLD is currently explained by a “multiple parallel-hit” hypothesis, which implicates the synergistic and concerted action of multiple events originating from various liver cell types (Tilg and Moschen, 2010).

Recent evidence indicates that the progression of MAFLD results from complex interactions within a gene–environment network, involving insulin resistance, mitochondrial dysfunction, oxidative and endoplasmic reticulum (ER) stress, gut dysbiosis, and alterations in the gut–liver axis and organokine signaling. These interconnected processes converge on chronic inflammation, ultimately driving the progression from MASH to cirrhosis and hepatocellular carcinoma (HCC) (Loomba et al., 2021a). Albeit these metabolic abnormalities are not the only ones, being a common nexus of other defects, they are considered to provide potential therapeutic targets to prevent or reverse liver inflammation and liver fibrosis. In keeping some drug candidates have been developed for the treatment of MAFLD and MASH by targeting hepatic lipid metabolism (Febbraio et al., 2019; Lazaridis and Tsochatzis, 2017; Smeuninx et al., 2020).

Some drug candidates have been developed for the treatment of MAFLD and MASH. Some of them target hepatic lipid metabolism and they have focused on pathways that affect the balance between fatty acid uptake and export, de novo lipogenesis (DNL), and fatty acid oxidation (FAO) in the liver (Febbraio et al., 2019; Lazaridis and Tsochatzis, 2017; Smeuninx et al., 2020). However, many have failed because of unwanted side effects (Chapman and Lynch, 2020; Kim et al., 2017). In March, 2024, resmetirom became the first US FDA-approved drug for the treatment of MASH and also approved for this indication by the EMA in August, 2025 (Li et al., 2026), nevertheless, until now with moderate effects.

Tumor suppressors, besides their actions regulating cell cycle and viability are emerging as important players mediating energy and glucose metabolism (Diehl et al., 2024; Fajas, 2013; Huber et al., 2021). Studies using mice deficient in cell cycle regulators have shown that the major phenotypes are metabolic perturbations. It is shown that some of these cell cycle regulators are crucial factors in metabolic control (Aguilar and Fajas, 2010), also in liver metabolism (Porteiro et al., 2017).

One of these cell cycle regulators is p107, a member of a pocket proteins (Rb family), which comprises pRb, p107 and p130 and their associated proteins. These proteins have been extensively studied in the relation to cell cycle control and tumorigenesis (Vidal et al., 2007) . The Rb family are implicated in the development and differentiation of many cell types and now are also known to have implications in adipogenesis and regulation of energy metabolism. p107 or Rb-like 1 (Rbl1) was identified through its interaction with SV40 Large T antigen and adenovirus E1S (Dyson et al., 1989; Ewen et al., 1991), p107 is a regulator of adipocyte differentiation (De Sousa et al., 2014; May et al., 2001; Richon et al., 1997) playing a key role in adipocyte lineage fates and the generation of pro-thermogenic adipocytes (Porras et al., 2017). Recently studies also demonstrated its implications in the regulation of energy metabolism (Hallenborg et al., 2009) (Cunarro et al., 2019) and more interesting, the total absence of p107, has been shown to improve hepatic metabolism in high-fat diet models (Cunarro et al., 2019).

Although studies studying p107 in liver are scarce, only seminal studies have connected p107 to growth arrest in neonatal hepatocytes (Timchenko et al., 1999), the fact that the Rb family act as repressors of the E2F family of transcription factors and they the finding that E2F is involved in liver regeneration (Ehmer et al., 2014) prompt us to assess the role of p107 on MAFLD and its progression to later stages of liver disease. In this context evidence regarding its actions in specific cell populations such hepatocytes or in the context of lipid metabolism and MALSD is scarce.

In the present study, we demonstrate that hepatic p107 expression is increased in patients and in preclinical models with MASLD. Using both global and liver-specific loss-of-function approaches, we show that p107 deficiency protects against hepatic steatosis through reduction of de novo lipogenesis and enhancement of mitochondrial activity. These findings are recapitulated in primary mouse and human hepatocytes. Conversely, p107 overexpression in vivo and in human hepatocytes promotes lipid accumulation. Unbiased phosphoproteomic analysis identifies fatty acid synthase (FASN) as a potential mediator of these effects. Together, these findings establish p107 as a regulator of hepatic lipid metabolism with important implications for the development of steatosis.

## 2. Material and methods

### 1. Cohort of patients

Liver biopsies were collected during bariatric surgery from a cohort of 23 patients with severe obesity at the Clínica Universidad de Navarra (Spain). For the purposes of this study, obesity was defined by a body mass index (BMI) ≥30 kg/m² and a body fat content ≥35%. BMI was determined using the standard formula (weight/height^2^), while total body fat was assessed via air-displacement plethysmography (Bod-Pod®, Life Measurements, Concord, CA, USA).

Inclusion criteria required a comprehensive diagnostic work-up, including physical examination, laboratory investigations, ultrasonography, and histological confirmation via liver biopsy. The diagnosis of MASH was established by an expert pathologist blinded to all experimental results, following the Kleiner and Brunt criteria (Kleiner et al., 2005). To evaluate disease severity, a MASLD activity score (NAS) ranging from 0 to 8 was determined by integrating histological features of steatosis, lobular inflammation, and hepatocyte ballooning. Exclusion criteria were: a) daily alcohol intake ≥20 g for women and ≥30 g for men; b) evidence of hepatitis B virus surface antigen or hepatitis C virus antibodies in the absence of a history of vaccination; c) use of drugs causing MASLD (i.e. amiodarone, valproate, tamoxifen, methotrexate, corticosteroids or anti-retrovirals); and d) presence of other specific liver diseases, such as autoimmune liver disease, haemochromatosis, Wilson’s disease, or α-1-antitrypsin deficiency.

Anthropometric, biochemical and clinical characteristics are described in Supplementary Table 1. All reported investigations were carried out in accordance with the principles of the Declaration of Helsinki, as revised in 2013, and approved by the Hospital s Ethical Committee responsible for research (protocol 2021.005). Written informed consent was obtained from all the participants.

### 2. Animals

Male wild type (WT) and p107 knock out (KO) mice (mixed-background C57BL6/129) were previously described (Lee et al., 1996; Vidal et al., 2007) and 8 weeks old male C57BL/6J mice; they were housed in air-conditioned rooms (22–24 °C) under a 12:12h light/dark cycle. Animal experiments were conducted using n = 7 – 11 independent animals per group. After weaning, the mice were fed with standard chow (STD) (#U8200G10R, SAFE Diets) or a high fat diet (HFD) (Research Diets D12,492;60% fat, 5.24 kcal g^−1^, Research Diets, New Brunswick, NJ) during specified times and experiments. Food intake and body weight were measured weekly during all experimental phases. Following the experimental protocols, animals were euthanized, livers were weighed and snap-frozen in dry ice, and all tissues were stored at –80°C until analysis. All procedures were approved by the Animal Care Research Bioethics Committee of the University of Santiago de Compostela (license 15012/2024/005) and conducted in accordance with Directive 2010/63/EU and the Spanish Royal Decree 53/2013.

### 3. Lentivirus design and production

To downregulate p107 in the liver, specific shRNA sequences targeting p107 or luciferase (control) were designed through the GPP Web Portal Tool (available at https://portals.broadinstitute.org/gpp/public/). The oligos targeting the transcripts of interest were synthesized and subcloned into pLKO.1 puro-GFP vectors (Addgene) as previously described (Covelo-Molares et al., 2022). In brief, HEK293T cells were cultured in high-glucose DMEM supplemented with 10% FBS, 2 mM l-glutamine, and 1% penicillin and streptomycin. Cells were plated at a density of 8 × 10^6^ cells per 150 mm dish and transfected 24 h later with PEI (polyethylenimine; 408727, Sigma–Aldrich), along with 20 μg of pLKO.shRNAs plasmids or p107 overexpression plasmids, and 10 μg of psPAX2 and pMD2.G packaging mix. The medium was replaced after 24 h, and virus-containing supernatants were collected 48 h and 72 h post-transfection. Lentiviral particles were concentrated using centrifugal filter units with a 0.22 μm pore size (UFC903024, Amicon). The target sequences of the shRNAs used in this study were as follows:

shRbl1 #1: GCTTGATGTTATCACCCTATA

shRbl1 #2: ATCTTTGCCAATGCTATAATG

shLuciferase: AACAGACAGTGCGTTTCAAATT

In addition, a lentiviral vector was utilized to overexpress p107, using a plasmid designed for sustained p107 expression from Vector Builder. EGFP-control lentivirus (VB160109-10005) and pLV-EGFP T2A:Puro-CMV-mRbl1 (VB190619-1030dhg).

### 4. Tail vein injections for in vivo lentiviral

Mice were held in a specific restrainer for intravenous injections, Tailveiner (TV-150, Bioseb). The injections into the tail veins were carried out using a 27 G x 3/8 (0.40 mm x 10 mm) syringe. Mice were administered with 100 μl of lentiviral Rbl1 shRNA (1x10^9^ TU/mL) or control shRNA (1x10^9^ TU /mL), 100 μl of lentiviral Rbl1 plasmid(1x10^9^ TU /mL) or control GFP (1x10^9^ TU /mL), diluted in saline. In the HFD cohorts, 7-week-old mice were injected with p107-shRNA lentivirus and then placed on the diet. Similarly, p107-KO mice received the p107 overexpression vector at 8 weeks of age before starting the HFD. Mice were euthanized and tissues were immediately snap-frozen on dry ice and stored at -80 °C until further analysis.

### 5. Isolation and culture of primary hepatocytes

Primary hepatocytes were isolated from male C57BL/6 WT mice via collagenase perfusion. In brief, animals were anesthetized with isoflurane, the abdomen was opened and a catheter was inserted into the inferior vena cava while the portal vein was cut. Next, liver was washed by perfusion with Krebs-Henseleit (KH) perfusion medium at 37 °C.

After the washing, EGTA 0.05% (w/v) was added to the KH medium, and the perfusion was maintained for 5 min. Finally, an enzymatic digestion was performed during 10-12 min with KH perfusion medium supplemented with Ca2+ and 50 mg/ml collagenase type I (LS004196, Worthington). After perfusion, the liver was gently disaggregated. The viable cells were purified by density centrifugation at 500xrpm for 5 min 3 times at 4 °C. Isolated pure hepatocytes were seeded over collagen-coated culture dishes at a density of 4 x 10^5^ cells/well in the medium for cell adhesion serum-free Dulbecco s modified Eagle s medium (DMEM), supplemented with 10% (v/v) FBS, 1% (v/v) Glutamine and 1% (v/v)Penicillin-Streptomycin solution. Experiments in primary hepatocytes were performed with n = 6 technical replicates derived from two independent hepatocyte isolations.

### 6. Human cell culture

THLE2 human hepatic cell line (American Type Culture Collection, ATCC) was cultured in bronchial epithelial cell basal medium (BEBM) supplemented with a growth factors BulleKit (Lonza/Clonetics Corporation), 70ng/mL phosphoethanolamine, 5 ng/mL epidermal growth factor, 10% (v/v) FBS and 1% (v/v) Glutamine-Penicillin-Streptomycin solution (MERCK). THLE-2 cells were grown on culture plates precoated with a mixture of 0.01 mg/ml fibronectin (#33010018, Sigma Aldrich, USA), 0.01 mg/ml bovine serum albumin (#A4503, Sigma Aldrich, USA) and 0.03 mg/ml collagen type I (#sc-136157, Santa Cruz, USA). The cell line was maintained at 37°C in a humidified atmosphere containing 5% CO2. Experiments performed in THLE2 cells were carried out with n = 6 technical replicates per condition.

### 7. p107 silencing and overexpression in vitro and cotransfection

3 × 10^5^ THLE2 cells were seeded in a six-well plate for the experiments. The cells were transfected with specific small-interference RNA (siRNA) to knockdown the expression of p107 (siRBL1, siGENOME ON-Target plus-SMART pool, Dharmacon, Cat# M-003298-02-0005). The control group was administered with a non-targeting siRNA (siControl, siGENOME ON-Target plus Non-Targeting pool, Dharmacon, Cat# D-001810-10-05). For the transfection in THLE2 cells we used Dharmafect 1 reagent (Dharmacon, T-2001): 6.5 µl of Dharmafect 1 diluted on 250 µl of optiMEM (Life Technologies, 31985070) mixed with 0.05 nmol of each siRNA diluted in 250µl of optiMEM; 500 µl of the transfection mixture was added into each well. For plasmid transfection, we used a DNA plasmid containing the sequence necessary to increase the expression of p107 (pCMV_neo_-p107HA) (Zhu et al., 1993) and an empty plasmid as control. For THLE2 cells we used Lipofectamine 2000 (Invitrogen, #11668-019): 4 µl Lipofectamine 2000 diluted in 250 µl of optiMEM mixed with 2.5 µg of DNA (p107 or empty control) diluted on 250 µl of optiMEM. Next, this mixture was added into each well and incubated for 6 h. The medium was replaced with fresh medium after 6 hours. Cells were collected after a total of 24 and/or 48 hours to check the efficiency of silencing or overexpression by Western blot.

1.5 × 10^5^ THLE2 cells were seeded in a twelve-well plate and transiently co-transfected with the p107 expression plasmid and a small interfering RNA targeting fatty acid synthase FASN (siFASN, siGENOME ON-Target plus-SMART pool, Dharmacon, Cat# M-003954-00-0005) using Lipofectamine 2000 following the manufacturer’s instructions. Briefly, for each well, two separate pre-mixtures were prepared: 2.5 μg of p107 plasmid and 0.05 nmol of siFASN were diluted in 250 μL of optiMEM, while 6 μL of Lipofectamine 2000 were diluted in an equal volume (250 μL) of optiMEM. After 5 minutes of incubation at room temperature, the solutions were combined and incubated for 20 minutes to allow the formation of transfection complexes. The resulting mixture (500 μL total) was then added dropwise to the cells. The medium was replaced with complete growth medium 6 hours post-transfection, and cells were harvested 24 hours later for Oil Red O staining and protein expression analysis.

### 8. Metabolic flux assays

The oxygen consumption rate (OCR) and extracellular acidification rate (ECAR) of THLE2 cells and primary mouse hepatocytes were measured at 37°C by high-resolution respirometry with the Seahorse Bioscience XFp Extracellular Flux Analyzer (AgilentTechnologies). Specifically, glycolytic flux in THLE2 cells was evaluated by monitoring ECAR levels. For the OCR and ECAR measurements, 1.5 × 10^4^ cells THLE2 cells and 1× 10^4^ primary mouse hepatocytes were seeded per well in a XFp cell culture microplate (103022-100, Seahorse Bioscience, Agilent Technologies). After 24 hours in culture or silencing/overexpression experiments, the medium of the cells was removed and replaced with pre-warmed assay medium, composed of Seahorse XF DMEM medium (103575100 Seahorse Bioscience, Agilent Technologies) containing 1 mM sodium pyruvate, 2 mM L-glutamine and 10 mM glucose, and cultured at 37°C in room air. After equilibration in assay medium for 1 hour, five basal measurements of OCR and ECAR were performed. Next, we sequentially added into cells wells the modulators of respiration oligomycin (Oligo, 1.5 µM), carbonyl cyanide-4 (trifluoromethoxy) phenylhydrazone (FCCP, 1 µM) and rotenone/antimycin (Rot/AA, 0.5 µM) (Cell Mito Stress Test Kit, 103010-100, Seahorse Bioscience, AgilentTechnologies) or the modulators rotenone/antimycin (Rot/AA) followed by 2-Deoxyglucose (2-DG, 50mM) to evaluate the glycolytic function (Glycolytic Rate Assay Kit, 103346-100, Seahorse Bioscience, Agilent Technologies). The following key parameters of mitochondrial function were calculated according to the manufacturer’s user guide (103011-400, Seahorse Bioscience, Agilent Technologies): basal respiration, ATP-linked respiration, proton leak, maximal respiration, spare capacity and nonmitochondrial oxygen consumption. In the same way, the next parameters of glycolytic function were calculated according to the manufacturer’s user guide: basal glycolysis, compensatory glycolysis and Post 2DG acidification (Glycolytic Rate Assay Kit). All results were normalized to protein content of each well.

### 9. Fatty acid oxidation in THLE2 cells

#### The rate of fatty acid oxidation

in cells was determined by measuring the amount of ^14^CO_2_ (complete oxidation) and the amount of ^14^C-acid-soluble metabolites (ASM) (incomplete oxidation) released from [1-^14^C]-palmitate oxidation, as described by others with minor (Hirschey et al., 2010; Vila-Brau et al., 2011). Briefly, cells were incubated for 4 hours with medium supplemented with 0.5% (w/v) fatty acid free BSA complexed with 0.2 mM palmitate containing 0.5 μCi/ml [1-^14^C]-palmitate (Perkin Elmer Inc). Medium was then collected in a tube containing Whatman filter paper soaked with 0.1 M NaOH in the cap and 500 μl of 3 M perchloric acid were added to release the CO_2_, which was captured in the filter paper. The acidified medium was centrifuged at 21,000 × g for 10 min to remove particulate matter. The radioactivity of CO_2_ captured by the filter papers and of ASM (the supernatants of the culture media) was measured by a scintillation counter and expressed relative to the cell protein.

#### FAO enzyme activity

was measured using a FAO Assay Kit (Biomedical Research Service Center, E-141) according to the manufacturer’s instructions. All samples were harvested with 100 µL 1×Cell Lysis Solution. 30 µL of each sample was added in duplicate to a 96-well plate on ice. Immediately, 40 µl control solution was added to one set of wells and 40 µl of reaction solution to the other set of wells. The contents were mixed by gentle swirling for 10 s. The plate was covered and incubated in a 37 °C humidified incubator for 120 min. Optical density was read at 492 nm (OD 492) using a microplate reader. The control well reading was subtracted from the reaction well reading for each sample and the subtracted OD represents the FAO activity of the sample. Enzyme activity was normalized by total protein concentration.

### 10. Histological procedures

*Hematoxylin-Eosin Staining.* Samples were fixed in 10% buffered formalin for 24 h and then dehydrated and embedded in paraffin. Sections of 3 µm thick-ness were made on a microtome and stained by the standard hematoxylin-eosin alcoholic procedure according to the manufacturer’s instructions (BioOptica, Italy). They were mounted with a permanent (non-alcohol, non-xylene based) mounting medium and evaluated and photographed by means of a BX51 microscope equipped with a DP70 digital camera (Olympus, Japan).

*Oil Red O Staining*. Frozen sections (8 µm thick) of the livers were made, fixed in 10% buffered formalin, and stained with filtered Oil Red O for 10 min. The sections were washed with distilled water, counter stained with Mayer’s hematoxylin for 3 min and mounted with an aqueous mounting (glycerin jelly). In all the histological staining, we used up to 4 representative microphotographs of each animal were taken at 20X or 40X with a BX51 microscope equipped with a DP70 digital camera (Olympus). Oil-Red O staining was imaged under identical acquisition parameters across all samples, including objective, magnification, illumination, intensity, exposure time and camera gain). Image quantification was performed using ImageJ software (National Institutes of Health and the Laboratory for Optical and Computational Instrumentation, LOCI, University of Wisconsin). For each image, the total Oil Red O-positive area was segmented by color-thresholding and quantified as integrated area. In parallel, the number of cells per field was determined with cell counter. Lipid content was expressed as stained area per cell by dividing the Red O-positive area by the correspond cell-count. These values were then normalized to the mean of the control group and reported as relative Oil Red O staining.

### 11. Gene Expression Analysis

Gene expression was assessed by real-time PCR (TaqMan, Applied Biosystems, USA), as previously described 24. Total RNA was isolated from liver and cells samples using TRIzol reagent (15596018, Invitrogen) according to the manufacturer’s instructions. 100 ng of total RNA were used for each RT reaction, and cDNA synthesis was performed using the SuperScript First-Strand Synthesis System (Invitrogen) and random primers. Negative control reactions, containing all reagents except the sample were used to ensure specificity of the PCR amplification. Real-time PCR was performed in duplicate using QuantStudio 5 (Applied Biosystems) and appropriate TaqMan assays (Applied Biosystems, Life Technologies). Cycling conditions: 50°C for 10 min, then 40 cycles of 95°C for 15 s and 60°C for 1 min. The specific TaqMan assays used are described in Supplementary Table 2. Expression levels were normalized to HPRT1 for each sample and the fold change value was determined from the equation 2^−ΔΔCt^.

### 12. Western blot analysis

Western blots were performed as previously described (Cunarro et al., 2019; Tovar et al., 2013). Briefly, total protein lysates from the liver (20 µg) or cell (6-8 µg) were subjected to SDS-PAGE, electrotransferred onto an Immuno-Blot polyvinylidene difluoridemembrane (Bio-Rad, USA), and probed with the antibodies indicated in Supplementary Table 3. The antibody concentration was 1:1000. For protein detection, horseradish peroxidase-conjugated secondary antibodies (Dako, Denmark) and chemiluminescence (Pierce ECL western blottingsubstrate, Thermo Scientific, USA) were used. X-ray film (Super RX, Fuji Medical X-Ray Film, Fujifilm, Japan) was then exposed to the membranes and developed with developer and fixing liquids (AGFA, Germany) under appropriate dark-room conditions. Densitometric quantification was performed using ImageJ 1.52p software, and protein levels were normalized to GAPDH for each sample and expressed relative to controls.

### 13. Proteomics and Phospho-Proteomics

Mouse livers from p107 knockout and shLucif control groups subjected to a high-fat diet were homogenized using a FASTPREP system in miST buffer and subsequently diluted with SP3 buffer. For each sample, 180 µg of protein was digested following the SP3 method using magnetic Sera-Mag Speedbeads (Cytiva 45152105050250, 50 mg/mL). After heating for 10 min at 75°C, cysteines were alkylated with 32 mM (final) iodoacetamide for 45 min at room temperature in the dark. Beads were added to the samples at a 10:1 (w:w) ratio, and proteins were precipitated with ethanol (60% final concentration). Following three washes with 80% ethanol, on-bead digestion was performed in 40 µL of 50 mM Hepes (pH 8.3) with 2 µg of trypsin (Promega #V5073). Peptides were labeled using a TMT multiplex kit and pooled into a single mixture. The pooled sample was divided for total proteome and phosphoproteome analyses. For the total proteome, the mix was fractionated into 6 Strong Cation Exchange (SCX) fractions. Phosphopeptide enrichment by IMAC was performed on the remaining TMT-labelled material using the High-Select™ Fe-NTA Phosphopeptide Enrichment Kit (Thermo Fisher Scientific, Prod. Number A32992) according to the manufacturer’s instructions. IMAC eluates were subsequently dried and resuspended for analysis in 2% acetonitrile and 0.05% trifluoroacetic acid (TFA).

Data-dependent LC-MS/MS analyses were carried out on a Fusion Tribrid Orbitrap mass spectrometer (Thermo Fisher Scientific) connected through a nanoelectrospray ion source to an Ultimate 3000 RSLCnano HPLC system (Dionex) via a FAIMS Pro interface. Peptides were separated on a reversed-phase nano-LC column using an acetonitrile gradient in 0.1% formic acid. The total proteome fractions were analyzed using 3 FAIMS methods per fraction (23 injections in total), while the IMAC-enriched sample was analyzed using 5 FAIMS methods (5 injections).

Raw data files were analyzed with MaxQuant (version 2.6.7.0) against the Mus musculus reference proteome. Cysteine carbamidomethylation and TMT labelling (at peptide N-termini and lysine side chains) were selected as fixed modifications, while methionine oxidation and protein N-terminal acetylation were specified as variable modifications. Both peptide and protein identifications were filtered at a 1% False Discovery Rate (FDR) against a decoy database built by reversing protein sequences. The Match Between Runs (MBR) feature and TMT correction factors were applied. Data were normalized by median subtraction. Following routine quality control, two outlier samples (TMT channels 6 and 9) were entirely excluded from downstream analyses to ensure statistical robustness.

Differential analysis and visualization were conducted in R (version 4.5.2). To visualize the global distribution of expression changes, volcano plots for both the total proteome and the phosphoproteome (using P-sites matched to total proteins) were generated using the ggplot2 and ggrepel packages. Subsequently, to generate targeted expression heatmaps, the total proteome dataset was filtered using two distinct approaches to capture both broad trends and high-confidence targets. A less restrictive filtering strategy isolated proteins based solely on high statistical significance (p-value < 0.001). In contrast, a more stringent strategy was applied to isolate a high-confidence core of differentially expressed proteins, requiring > 3 combined razor and unique peptides, a p-value < 0.001, and an absolute Student’s t-test difference of means > 0.58. Proteins from both filtering tiers were mapped to specific murine Biological Process Gene Ontology (GO:BP) terms retrieved via the msigdbr and org.Mm.eg.db packages, and their expression differences were visualized as faceted heatmaps using ggplot2.

### 14. Data analysis and statistics

Data are reported as mean ± standard error of the mean (SEM). Normality was assessed using the Anderson–Darling test, and homoscedasticity was evaluated with the Fligner–Killeen test. For comparisons across treatment groups, Statistical significance was determined by two-tailed Student’s t-test or Mann-Whitney test when two groups were compared. For more that 2 groups one-way analysis of variance (ANOVA) was applied when model assumptions were met; otherwise, the Kruskal–Wallis rank-sum test was used. For longitudinal weight measurements, repeated-measures ANOVA or the Friedman rank-sum test (when normality was not satisfied) was conducted. Multiple comparisons were adjusted using the Holm method. On All statistical tests P value < 0.05 was considered statistically significant. Analyses were performed in R version 4.3.2 (R Core Team, 2023) using the statsplot package (Patil, 2021) or *GraphPad Prism version 9.0 (GraphPad Software, San Diego, CA, USA)*.

## 3. Results

### 2.1 p107 is increased in the liver of patients with MASLD/MASH

To explore the potential involvement of p107 in MASLD, we first quantified hepatic p107 mRNA levels in obese patients with or without MASLD and observed a significant increase in p107 expression in patients with MASLD (Fig. 1A). We then examined the association between p107 expression and disease severity, revealing a positive correlation with MASLD activity score (NAS) and fibrosis score (Fig. 1B) as well as serum triglyceride content (Fig. 1C). Collectively, these findings indicate that hepatic p107 expression is elevated in MASLD and is associated with disease progression towards MASH.

**Figure 1.**
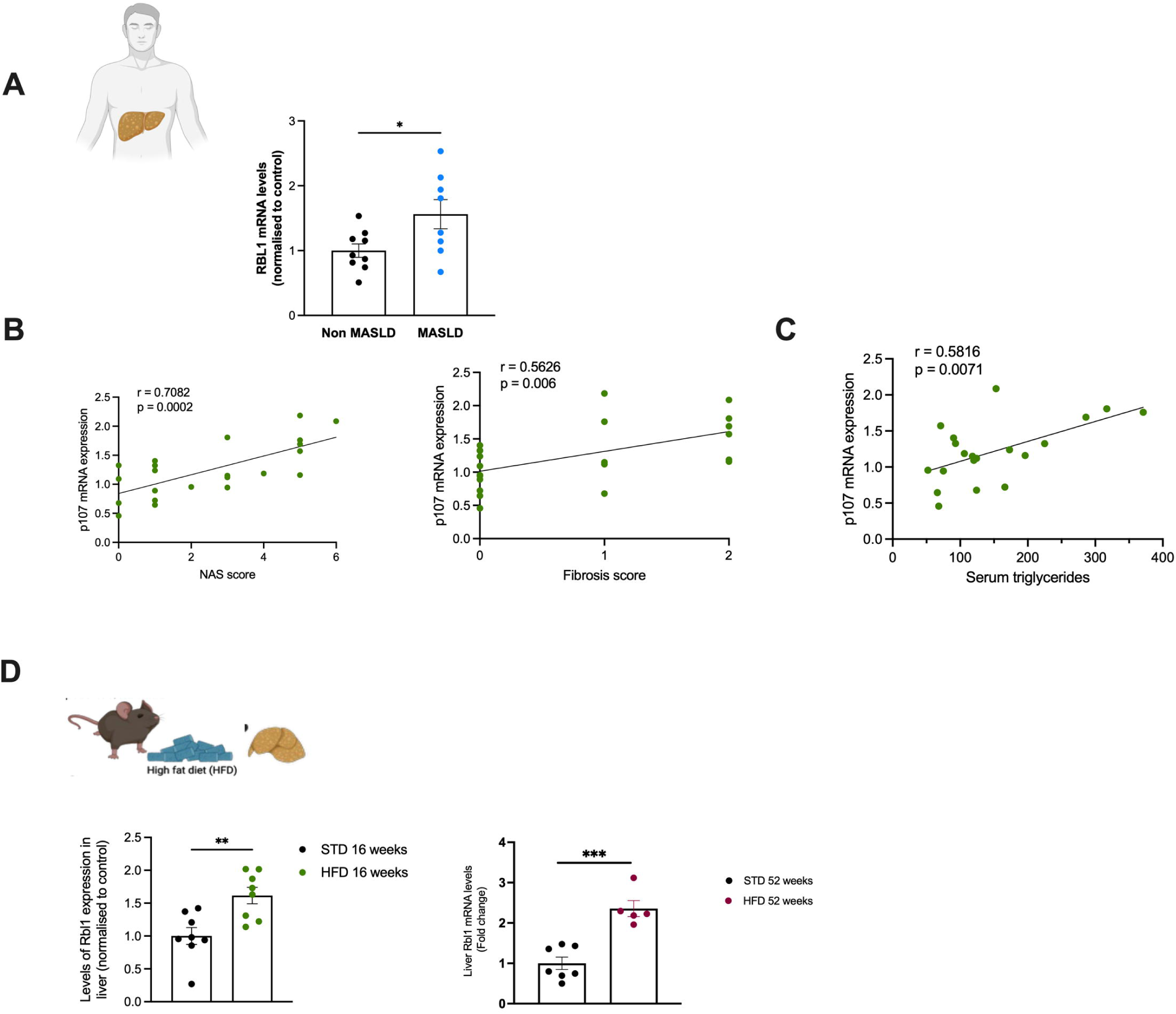
Liver p107 increase in patients with MASLD and it positively correlates with MASLD progression. (A) mRNA expression of p107 in liver of human patient with different stages of MASLD. (B) Correlation of human patient samples between hepatic p107 and NAS score and fibrosis score. (C) Correlation of p107 and triglycerides content. Pearson’s correlation test if the data follow a normal distribution or Spearman’s correlation test if the data do not follow a normal distribution. (D) mRNA expression of p107 in mice feed with SDT diet or HFD during 16 weeks (left panel) and 53 weeks (right panel). HPRT was used to normalize mRNA levels. Data are expressed as mean ±SEM. *p < 0.05, **p < 0.01, ***p <0.001, using a Student’s *t test*.

### 2.2. p107 deficiency prevents HFD-induced liver steatosis, inhibits de novo lipogenesis and increases fatty acid oxidation in liver

We next investigated the role of p107 in preclinical models of MASLD. We first assessed hepatic p107 expression in mouse models of diet-induced steatosis, including mice fed a high-fat diet (HFD) for 16 weeks and 52 weeks to model more advanced disease (Fig. 1D, left and right panels, respectively). In both settings, p107 expression was increased, indicating that its upregulation is associated with steatosis progression in preclinical models.

To determine the functional relevance of p107 *in vivo*, we subjected global p107 knockout (p107KO) mice to HFD feeding for 16 weeks. p107KO mice exhibited reduced weight gain compared with control animals (Fig. 2A), consistent with previous findings from our group (Cunarro et al., 2019). Moreover, p107KO mice were protected from hepatic steatosis, as evidenced by Oil Red O staining (Fig. 2B). Similar results were observed in female, which showed reduced weight gain and protection against liver steatosis (Supplementary Fig. 1E, F), indicating that this protective effect is sex-independent.

**Figure 2.**
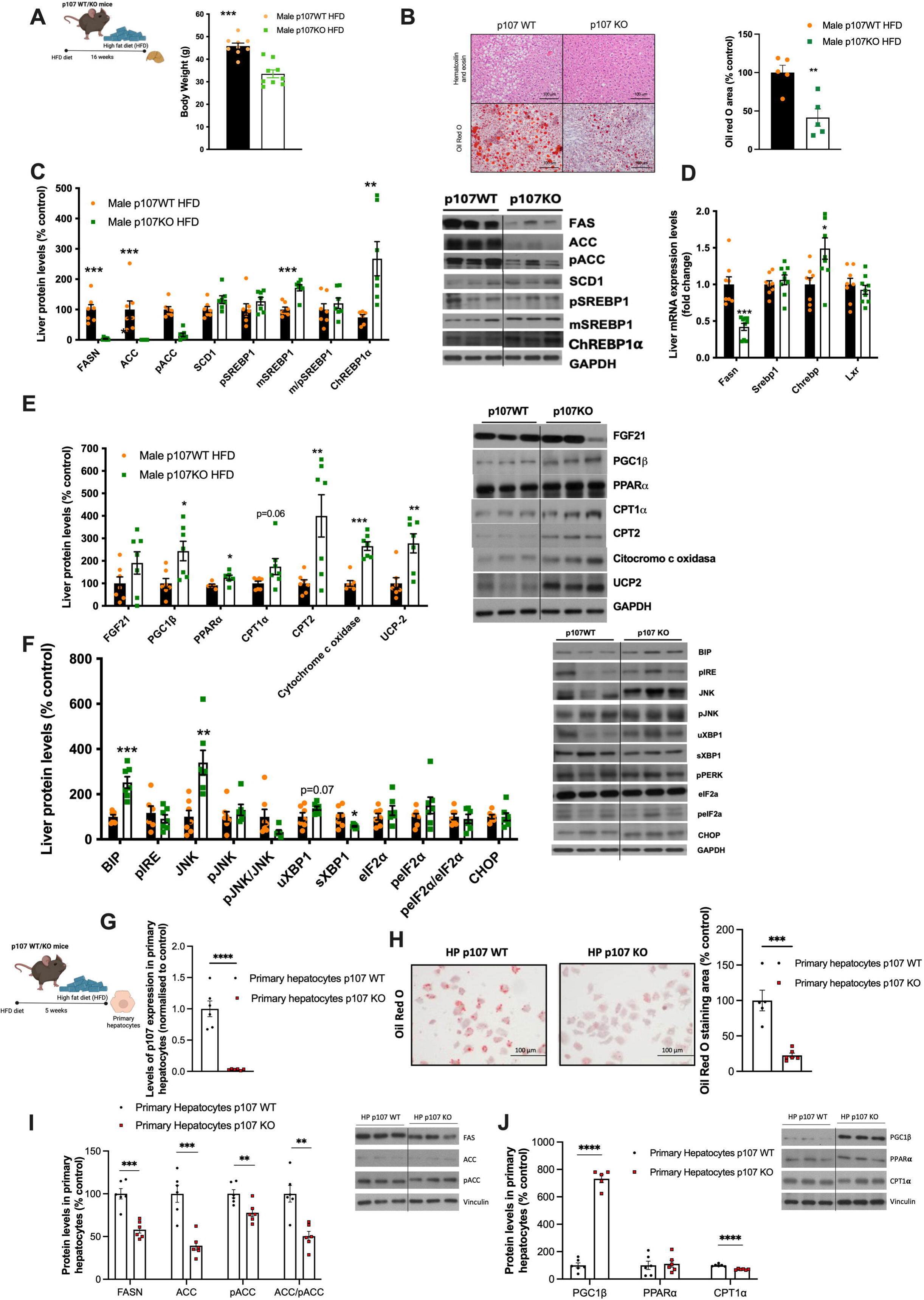
p107 KO mice are resistant to steatosis under very high-fat diet (HFD) and p107 regulates lipid accumulation in primary hepatocytes. p107 KO and control mice were fed a HFD for 16 weeks. (A) Body weight of control mice (n = 9) and p107 KO mice (n = 10). (B) Representative microphotographs of H&E staining (upper pannel) and Oil Red O (lower pannel) and Oil Red O semiquantification (right pannel) of liver sections (n=5-6). (C) Quantification of immunoblot analysis of *de novo lipogenesis* markers in liver (n = 7 per group) and a representative immunoblot. (D) mRNA expression of *de novo lipogenesis* markers in liver (n = 8 per group). (E) Quantification of immunoblot analysis of lipolysis and mitochondrial activity markers in liver (n = 7 per group) and a representative immunoblot. (F) Quantification of immunoblot analysis of ER stress markers in liver (n = 7 per group) and a representative immunoblot. HPRT was used to normalize mRNA levels, and GAPDH was used to normalize protein levels. (G) p107 protein levels in primary hepatocytes from p107KO and WT mice under HFD (n=6 per group). (H) Representative microphotographs and quantification of Oil Red O staining from primary hepatocytes for 24 hours (n = 6 per group). (I) Quantification of immunoblot analysis of *de novo lipogenesis* markers (n = 6 per group) and a representative immunoblot in hepatocytes for WT and p107KO mice under HFD. (J) Quantification of immunoblot analysis β-oxidation markers (n = 3 per group) and a representative immunoblot in hepatocytes for WT and p107KO mice under HFD Data are expressed as mean ±SEM. *p < 0.05, **p < 0.01, ***p <0.001, using a Student’s *t test*.

P107KO mice exhibited a marked reduction in the expression of key proteins involved in *de novo lipogenesis* (DNL), including FAS (fatty acid synthase) and ACC (acetyl-CoA carboxylase), in both males (Fig. 2C) and females (Supplementary Fig. 1H). The decrease in FAS was further confirmed at the mRNA level (Fig. 2D). In contrast, the expression of proteins involved in mitochondrial fatty acid β-oxidation, such as CPT1α and CPT2, was increased, together with elevated levels of the transcription factor PPARα (Fig. 2E and Supplementary Fig. 1E). Consistently, markers of mitochondrial activity, including mitochondrial complex IV (cytochrome c oxidase) and UCP2, were also upregulated (Fig. 2E).

HFD-fed livers are typically exposed to metabolic stress associated with lipid accumulation, leading to activation of the unfolded protein response (UPR). Notably, p107KO mice showed increased expression of the ER chaperone BiP (GRP78), a key regulator of protein folding and ER homeostasis (Fig. 2F), without significant changes in other ER stress markers. These findings suggest that p107 deficiency may preserve the correct folding of the proteins, thereby preventing the activation of a full UPR.

### 2.3 Primary hepatocytes from p107KO mice are protected from high fat-induced lipid accumulation and display reduced de novo lipogenesis and enhanced oxidative capacity

To determine whether the effects of p107 are directly in hepatocyte, we first quantified p107 expression in primary hepatocytes isolated from mice fed a standard diet (STD) or a high-fat diet (HFD) for 11 or 16 weeks. p107 expression was significantly increased in hepatocytes in a time-dependent manner upon HFD feeding (Supplementary Fig. 1F), indicating that its upregulation occurs directly in hepatocytes and is not secondary to systemic alterations in other tissues such as adipose tissue.

We next assessed lipid accumulation and metabolic pathways in primary hepatocytes isolated from WT and p107KO mice under HFD conditions. As expected, p107 expression was undetectable in p107KO hepatocytes (Fig. 2G). Notably, p107KO hepatocytes exhibited reduced lipid accumulation compared with WT cells, as determined by Oil Red O staining (Fig. 2H). Consistent with the in vivo findings, p107 deficiency resulted in decreased expression of key lipogenic proteins, including FASN, ACC and pACC (Fig. 2I). In parallel, the expression of the oxidative regulator PGC1β was increased (Fig. 2J), suggesting enhanced mitochondrial lipid oxidation.

### 2.4 Liver-specific downregulation of p107 protects against HFD-induced steatosis

To determine whether the protective effects observed in global p107KO mice are hepatocyte-autonomous and independent of systemic factors such as adipose tissue function or thermogenesis, we generated a liver-specific p107 knockdown model using virogenetic delivery of shRNA targeting p107. This approach resulted in a marked reduction of hepatic p107 expression (Fig. 3A). Following viral delivery, mice were fed a high-fat diet (HFD) for 11 weeks. Liver-specific p107 knockdown mice displayed comparable body weight gain to control animals (Fig. 3B). However, histological analysis and Oil Red O staining revealed a clear reduction in hepatic lipid accumulation in p107-deficient livers (Fig. 3C), recapitulating the phenotype observed in the liver of global p107KO mice.

**Figure 3.**
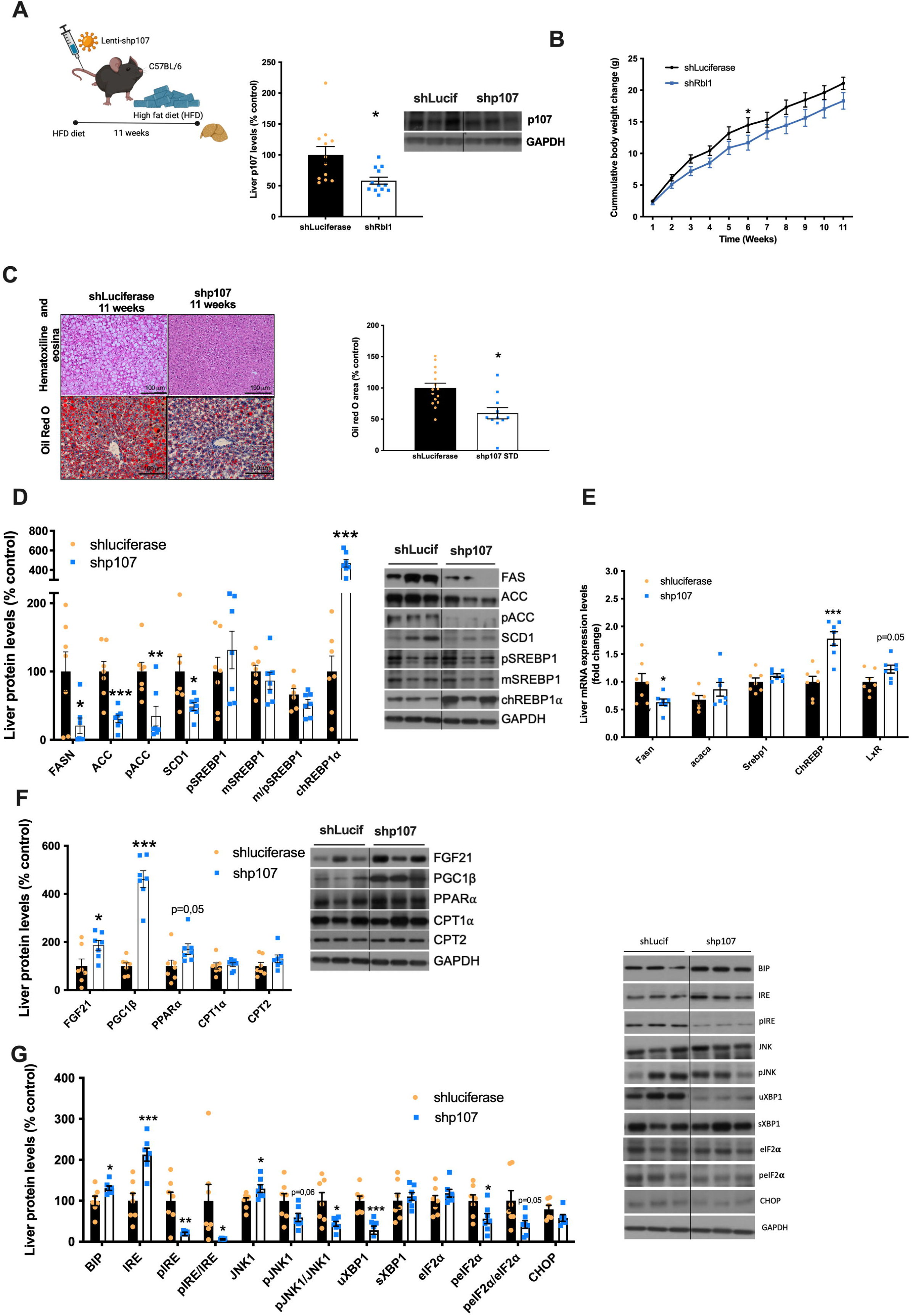
Silencing of hepatic p107 in mice fed a very high-fat diet (HFD) prevents the development of steatosis. Mice receiving the tail vein injection of lentivirus encoding shRNA p107 or luciferase were fed a HFD for 11 weeks. (A) Quantification of immunoblot analysis of expression of p107 in liver after shRNA-Luciferase administration (n = 12) or shRNA-p107 (n = 12) and a representative immunoblot. (B) Cumulative body weight change. (C) Representative microphotographs of H&E staining (upper panel) and Oil Red O (lower panel) and semiquantification (right panel) of liver sections. Oil Red O staining was quantified using ImageJ (n = 12-15 per group). (D) Quantification of immunoblot analysis of *de novo lipogenesis* markers (n = 7 per group) and a representative immunoblot. (E) mRNA expression of *de novo lipogenesis* markers (n = 7 per group). (F) Quantification of immunoblot analysis of β-oxidation markers (n = 7 per group) and a representative immunoblot. (G) Quantification of immunoblot analysis of ER stress markers in liver (n = 7 per group) and a representative immunoblot. HPRT was used to normalize mRNA levels, and GAPDH was used to normalize protein levels. Data are expressed as mean ±SEM. *p < 0.05, **p < 0.01, ***p <0.001, using a Student’s *t test*.

Consistent with these findings, hepatic p107 downregulation led to decreased protein levels of key de novo lipogenesis (DNL) markers, including FASN, ACC and SCD1 (Fig. 3D). The reduction in FASN was further confirmed at the mRNA level (Fig. 3E). In parallel, p107 knockdown increased the protein levels of regulators of lipid oxidation, including FGF21, PGC1β and PPARα, whereas no significant changes were observed in the fatty acid transporters CPT1α and CPT2 (Fig. 3F).

Importantly, liver-specific p107 inhibition was associated with reduced activation of ER stress signaling pathways, as indicated by decreased phosphorylation ratios of IRE1α, JNK and eIF2α. In addition, increased protein levels of the ER chaperone BiP (GRP78) was observed (Fig. 3G), consistent with improved protein folding capacity and reduced ER stress.

### 2.5 Hepatic inhibition of p107 primarily affects lipid accumulation prior to ER stress

To delineate the temporal sequence of events, we performed a shorter HFD intervention (7 weeks) to determine whether alterations in lipid accumulation precede changes in ER stress. Efficient hepatic knockdown of p107 was confirmed (Supplementary Fig. 2A), with no significant differences in body weight between groups (Supplementary Fig. 2B). Despite this, liver-specific p107 deficiency resulted in reduced hepatic lipid accumulation, as assessed by histological analysis and Oil Red O quantification (Supplementary Fig. 2C).

At this early time point, p107-deficient livers exhibited a marked decrease in FASN expression (Supplementary Fig. 2D) together with increased levels of PGC1β (Supplementary Fig. 2E), indicating early suppression of *de novo* lipogenesis and activation of oxidative pathways. In contrast, no significant changes were observed in ER stress markers (Supplementary Fig. 2F). These findings suggest that inhibition of de novo lipogenesis represents the primary event, leading to reduced lipid accumulation and subsequently preventing the development of ER stress.

### 2.6 Hepatic restoration of p107 reverses the protective metabolic phenotype in p107KO mice

To assess whether re-expression of p107 in the liver is sufficient to reverse the metabolic phenotype observed in p107KO mice, we performed a gain-of-function approach by delivering a lentiviral vector encoding p107 via tail vein injection in HFD-fed p107KO mice (Fig. 4A). Hepatic restoration of p107 expression did not affect body weight (Fig. 4B). However, it markedly increased hepatic lipid accumulation (Fig. 4C), reversing the steatosis-resistant phenotype observed in p107KO mice.

**Figure 4.**
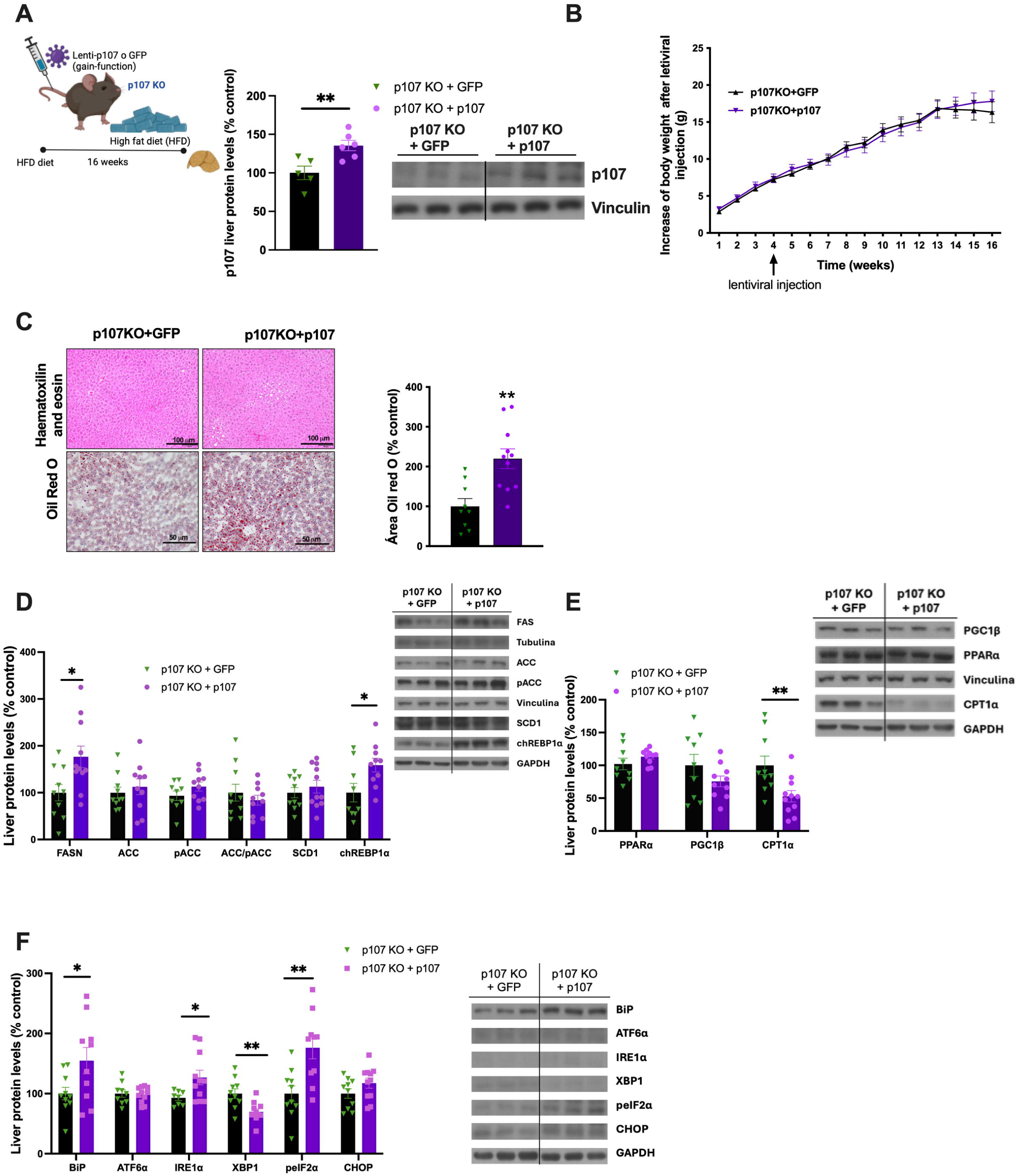
Recovery of hepatic p107 expression in the liver of p107 ko mice reverses the effects on hepatic fat accumulation. p107 KO mice were fed a HFD for 16 weeks, received the tail vein injection of lentivirus encoding p107 or GFP in week 4 of the diet. (A) Quantification of immunoblot analysis of expression of p107 in liver p107 KO + GFP (n = 6) and in liver p107 KO + p107 (n =6) and a representative immunoblot. (B) Cumulative body weight change. (C) Representative microphotographs of H&E staining (upper panel) and Oil Red O (lower panel) of liver sections. Oil Red O staining was quantified using ImageJ (n = 9-11 per group). (D) Quantification of immunoblot analysis of *de novo lipogenesis* markers (n = 9-11 per group) and a representative immunoblot. (E) Quantification of immunoblot analysis of β-oxidation markers (n = 9-11 per group) and a representative immunoblot. (F) Quantification of immunoblot analysis of ER stress markers in liver (n = 9-11 per group) and a representative immunoblot. GAPDH, 𝛂-tubulin and vinculin were used to normalize protein levels. Data are expressed as mean ±SEM. *p < 0.05, **p < 0.01, ***p <0.001, using a Student’s *t test*.

Consistently, re-expression of p107 restored the protein levels of the lipogenic enzyme FASN (Fig. 4D), indicating reactivation of de novo lipogenesis. In parallel, expression of the mitochondrial fatty acid transporter CPT1α was reduced (Fig. 4E), suggesting a shift away from lipid oxidation.

Importantly, hepatic p107 restoration was associated with increased activation of ER stress signaling, as reflected by elevated phosphorylation of eIF2α (Fig. 4F), further supporting a link between p107 activity, lipid accumulation and ER stress.

### 2.7 Inhibition of p107 in human hepatocytes reduces lipid accumulation by suppressing de novo lipogenesis and enhancing lipid oxidation

To further investigate the mechanisms underlying p107 function and assess its relevance in human systems, we used the human hepatocyte cell line THLE-2. p107 expression was efficiently silenced using small interfering RNA (siRNA), as confirmed 48 hours post-transfection (Fig. 5A). p107 knockdown markedly reduced intracellular lipid accumulation compared with control cells, as assessed by Oil Red O staining (Fig. 5B). At the molecular level, p107 silencing led to a clear reduction in the expression of key lipogenic markers, including FASN, at both protein and mRNA levels (Fig. 5C, D). In addition levels of SCD1, a key enzyme involved in fatty acid desaturation, was also decreased (Fig. 5C). Consistent with a global suppression of de novo lipogenesis (DNL), metabolic flux analysis using radiolabeled acetate revealed a significant reduction in the de novo synthesis of free fatty acids, with a trend towards decreased triglyceride synthesis (Fig. 5E).

**Figure 5.**
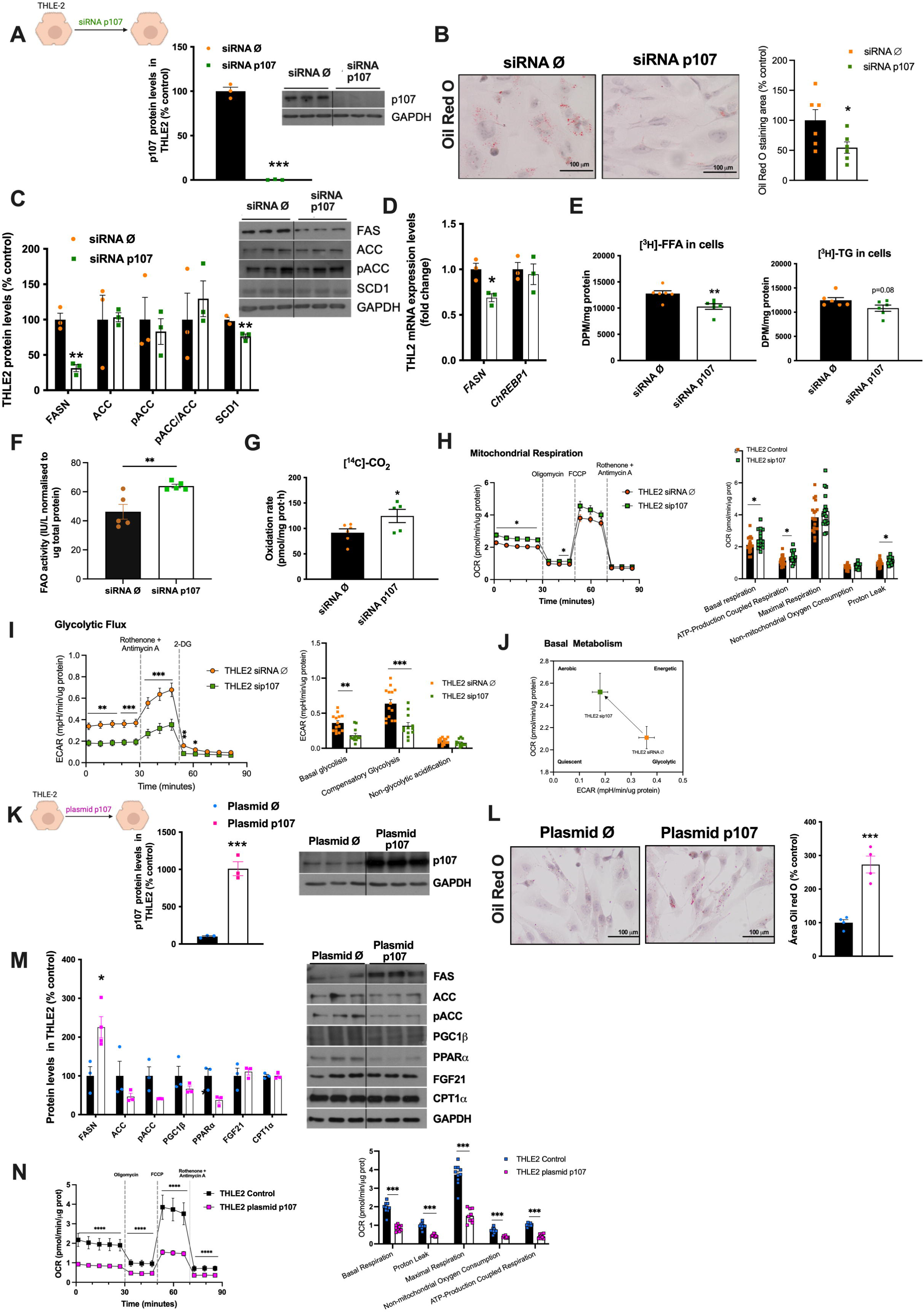
p107 regulates lipid accumulation in human hepatic cell line (THLE2). (A) p107 protein levels in THLE2 cells transfected with siRNA p107 or siRNA control for 48 hours (n=3 per group). (B) Representative microphotographs of Oil Red O staining (left pannel) of THLE2 cells downregulating p107 (sip107) for 48 hours (n = 3 per group) and Oil Red O semiquantification (right pannel) (n = 3 per group). (C) Quantification of immunoblot analysis of *de novo lipogenesis* markers (n = 3 per group) and a representative immunoblot. (D) RNA expression of *de novo lipogenesis* markers (n = 3 per group. (E) *De novo* synthesis of free fatty acids (FFA) and triglycerides (TG) in THLE2 (n = 6 per group). (F) FAO activity in THLE2 cells transfected with siRNA p107 or siRNA control for 48 hours (n=5 per group). (G) Oxidation rate of palmitic acid (n = 6 per group). (H and I)) OCR and (I) ECAR of p107-silenced THLE2 (n = 17-19 per group). (J) Basal energetic metabolic states, based on quantification of ECAR and OCR during basal metabolism. (K) p107 protein levels in THLE2 after overexpressing p107 (n = 3 per group). (L) Representative microphotographs (left pannel) and semiquantification (right pannel) of Oil Red O staining of THLE2 cells overexpressing p107 (plasmid p107) for 24 hours. Oil Red O staining was quantified using ImageJ and normalized to the total number of nuclei per field (n = 4 per group). (M) Quantification of immunoblot analysis of *de novo lipogenesis* markers (n = 3 per group) and a representative immunoblot. (N) OCR of p107-overexpressed THLE2 (n =10 per group). GAPDH was used to normalize protein levels. Data are expressed as mean ±SEM. *p < 0.05, **p < 0.01, ***p <0.001, using a Student’s *t test*.

In contrast, p107 inhibition enhanced lipid oxidative pathways. Fatty acid oxidation (FAO) activity was increased (Fig. 5F), and analysis of β-oxidation rates using ¹⁴C-palmitate confirmed enhanced lipid oxidation in p107-silenced cells (Fig. 5G). Moreover, mitochondrial function was improved, as indicated by increased oxygen consumption rate (OCR), including basal respiration, ATP-linked respiration and mitochondrial capacity (Fig. 5H). In parallel, glycolytic flux was reduced (Fig. 5I), suggesting a metabolic shift towards oxidative metabolism.

Together, these findings indicate that p107 downregulation promotes an aerobic basal metabolism (Fig 5J).

### 2.8 Overexpression of p107 promotes lipid accumulation and impairs mitochondrial function in human hepatocytes

To complement the loss-of-function studies, we assessed the effects of p107 overexpression in THLE-2 cells. Efficient overexpression of p107 was achieved following plasmid transfection (Fig. 5K). In contrast to p107 silencing, p107 overexpression led to a marked increase in intracellular lipid accumulation, as determined by Oil Red O staining (Fig. 5L).

At the molecular level, p107 overexpression resulted in increased expression of the lipogenic enzyme FASN and reduced levels of the oxidative regulator PPARα (Fig. 5M), indicating a shift towards lipid storage. Consistently, mitochondrial function was impaired, as evidenced by decreased oxygen consumption rate (OCR), including both basal respiration and maximal respiratory capacity (Fig. 5N).

### 2.9 Phosphoproteomic profiling links p107 to lipid metabolic pathways

To identify signaling pathways through which p107 regulates hepatic lipid metabolism, we performed large-scale proteomic and phosphoproteomic analyses in livers from WT and liver-specific p107-deficient mice under HFD conditions. Volcano plot analysis revealed widespread changes in both protein abundance (Fig. 6A) and phosphorylation sites (Fig. 6B) upon p107 knockdown, and most of these were showing a downregulation pattern. In a heatmap grouped by GO Biological Process terms and filtered by significance we found expressed proteins highlighted a strong enrichment of lipid metabolism-related pathways (Fig 6C). Subsequent utilizing strict filtering of these datasets identified FASN as a prominent candidate (Fig. 6D) potentially mediating the effects of p107 on hepatic lipid accumulation. These data provide further mechanistic support for the role of p107 in regulating de novo lipogenesis through lipid metabolic networks.

**Figure 6.**
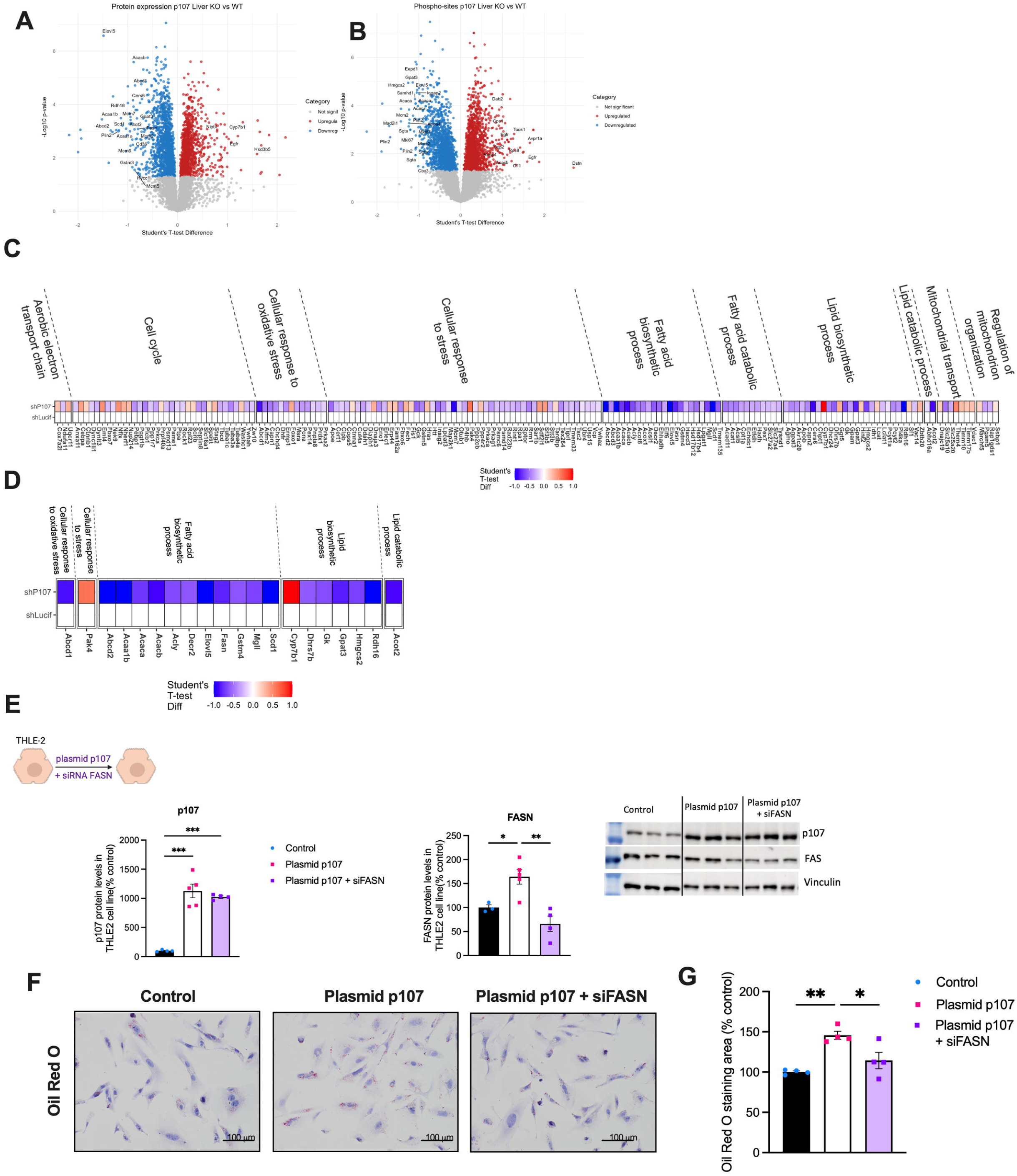
p107 is dependent of FAS in the effects on hepatic fat accumulation. (A) Volcano plot of total protein expression in p107 liver KO mice compared to shLucif controls (n = 7). Red and blue points indicate significantly up- and downregulated proteins (p < 0.05). (B) Volcano plot of phosphosite abundance in p107 liver KO mice compared to shLucif controls (n = 7). Points represent individual phosphosites—annotated by their parent protein name—with red and blue indicating significant changes (p < 0.05). (C) Heatmap of total protein expression differences grouped by GO Biological Process terms, filtered by significance (p < 0.001). (D) Heatmap of robust total protein expression differences grouped by GO terms, utilizing strict filtering (> 3 combined razor and unique peptides, p < 0.001, absolute t-test difference > 0.58). (E) Quantification of p107 (left panel) and FAS (right panel) in THLE2 after overexpressing p107 (plasmid p107) and silencing FAS (siRNA FAS) during 24h (n = 3 per group) and representative immunoblot. (F) Representative microphotographs of Oil Red O staining of THLE2 cells overexpressing p107 (plasmid p107) and silencing FAS (siRNA FAS) for 24 hours. (G) Semiquantification of Oil Red O staining. Vinculin was used to normalize protein levels. Data are expressed as mean ±SEM. *p < 0.05, **p < 0.01, ***p <0.001, using a Student’s *t test*.

### 2.10 FASN mediates the effects of p107 on hepatocellular lipid accumulation

Based on the proteomic data indicating alterations in key enzymes of de novo lipogenesis (DNL), together with our in vivo and in vitro findings, we next investigated whether FASN mediates the effects of p107 on lipid accumulation. To test this, we performed co-transfection experiments in THLE-2 cells combining FASN siRNA with p107 overexpression. Cells were first transfected with FASN siRNA to achieve efficient knockdown, followed by transfection with an p107-expressing plasmid, which increased p107 levels while maintaining FASN suppression (Fig. 6D, E).

As expected, p107 overexpression alone increased intracellular lipid accumulation, as assessed by Oil Red O staining. However, co-transfection with FASN siRNA markedly attenuated this effect, significantly reducing lipid accumulation compared with p107 overexpression alone (Fig. 6G). These results indicate that, the effects of p107 in lipid accumulation are, at least in part, dependent on FASN activity, supporting a key role for the DNL pathway in mediating p107-driven lipid accumulation.

## 4. Discussion

In this study, we demonstrated for the first time the role of p107 as a regulator of hepatic lipid metabolism. We generated a rodent model with liver-specific inhibition of p107, which allowed us to demonstrate that the absence of p107 in the liver protects mice fed a high-fat diet (HFD) from the development of diet-induced steatosis highlighting a hepatocyte-autonomous role in metabolic adaptation. These findings are supported by complementary data in primary mouse hepatocytes, human hepatic cell lines, and patients with MASLD, underscoring the translational relevance.

p107 deficiency induces a coordinated metabolic rewiring characterized by suppression of de novo lipogenesis (DNL) and enhanced mitochondrial lipid oxidation. Consistently, p107-deficient livers display reduced expression of key lipogenic regulator, including FASN and ACC (acetyl-CoA carboxylase), together with increased expression of genes involved in fatty acid oxidation. Given that hepatic lipid accumulation in MASLD is driven by increased DNL (Ipsen et al., 2018; Lawitz et al., 2022) and impaired oxidation (Loomba et al., 2021a), this shift towards lipid utilization likely underlies the reduced hepatic lipid burden observed in our model. This is in concordance with other mice models where the reduction of DNL (Kim et al., 2017) or increase in fatty acid oxidation (Cho et al., 2017) alleviated MASLD. In this context, our findings provide a framework for translational efforts aimed at developing therapeutic strategies to decrease hepatic DNL by inhibiting ACC and FAS, has been already applied (Harriman et al., 2016; Loomba et al., 2018; Loomba et al., 2021b).

In parallel, p107 deficiency attenuates ER stress, as evidenced by reduced activation of unfolded protein response (UPR) pathways and increased expression of the chaperone BIP. Given the well-established bidirectional relationship between ER stress and lipid metabolism (Lee et al., 2012; Zheng et al., 2010), our data support a model in which suppression of DNL represents the initiating event. Time-course experiments indicate that inhibition of FAS-dependent lipogenesis precedes resistance to ER stress, whereas increased mitochondrial lipid oxidation emerges only after prolonged exposure, suggesting that enhanced oxidation represents a secondary adaptive response. These effects are recapitulated in cell-autonomous systems. In both human hepatic cell lines and primary mouse hepatocytes, inhibition of p107 reduces lipid accumulation, indicating a direct role in hepatocellular lipid handling and increase fatty acid oxidation in concordance with other preclinical studies (Gonzalez-Rellan et al., 2023; Patitucci et al., 2023). Also, the OCR and ECAR indicate p107 downregulation promotes mitochondrial activity, translated into a healthy hepatic metabolism. Mechanistically, proteomic analyses and functional experiments identify FASN-driven DNL as a key downstream effector of p107. Overexpression of p107 enhances lipid accumulation and promotes a lipogenic program, whereas silencing of FASN attenuates these effects, supporting a model in which p107 acts upstream of the FASN axis. These findings indicate that the pro-steatotic effects of p107 are, at least in part, dependent on FASN activity, consistent with its established role in hepatic steatosis. However, the relationship between FASN and the improvement of hepatic steatosis appears to be context dependent. Notably, liver-specific genetic ablation of FASN in adult mice has been reported to produce minimal metabolic alterations under standard dietary conditions, with no significant changes in hepatic lipid content (Chakravarthy et al., 2005). In the other end, these animals develop marked steatosis when exposed to a fat-free diet. Despite these observations, a substantial body of preclinical evidence supports a pathogenic role for FASN in MASLD (O’Farrell et al., 2022). Moreover, ex vivo studies in human hepatic stellate cells indicate that FASN inhibition attenuates fibrogenesis, further supporting its role in disease progression (Bates et al., 2020; Wei et al., 2016).

Consistent with this model, genetic restoration of p107 expression in the liver of p107-deficient mice fed an HFD reversed both the resistance to hepatic lipid accumulation and the suppression of FAS expression thus adding further support to the notion that liver-specific inhibition of p107 confers protection against diet-induced steatosis primarily through the downregulation of DNL and modulation of lipid oxidation and attenuation of ER stress (Lee et al., 2012; Zheng et al., 2010).This is particularly relevant for several reasons. We previously reported that a KO of p107 prevents diet-induced obesity by increasing energy expenditure via increased thermogenesis in BAT and browning of WAT (Cunarro et al., 2019), suggesting that organ crosstalk mechanisms could be influencing liver steatosis. These findings indicate that hepatic p107 is sufficient to regulate lipogenic processes in the liver and suggest a role in metabolic flux control independent of its function in cell cycle regulation, extending beyond E2F-dependent pathways.

From a broader perspective, our results extend the emerging paradigm that cell cycle regulators actively participate in metabolic control (Fondevila et al., 2024; Gonzalez-Rellan et al., 2021; Lopez-Mejia et al., 2017) increasing evidence supports a bidirectional relationship between cell cycle machinery and metabolism. By simultaneously suppressing FASN–mediated lipogenesis, attenuating ER stress, and promoting mitochondrial lipid oxidation, inhibition of p107 targets multiple key mechanisms underlying MASLD pathogenesis. In this context, p107 may act at the interface between proliferative signaling and metabolic control, potentially integrating cues from pathways such as CDK–cyclin complexes (Chibazakura et al., 2004), E2F transcription factors (Burkhart et al., 2010), and nutrient-sensing pathways (Cuyas et al., 2014; Zhang et al., 2025). The conservation of these effects in human hepatocytes, together with their dependence on FASN, further strengthens the translational potential of this pathway.

Several limitations should be acknowledged. Although our data supports a central role for the FASN axis, additional downstream pathways—including lipid droplet dynamics, autophagy (lipophagy), and mitochondrial function—may also contribute to the observed phenotype. Given the role of p107 as a transcriptional corepressor (Nicolas et al., 2003; Scime et al., 2005), it will be important to determine whether p107 directly binds regulatory regions of the FASN gene or instead modulate DNL indirectly (De Sousa et al., 2014; Nicolas et al., 2003; Scime et al., 2010). A plausible model is that these effects are mediated through transcription factors such as E2F, since it was described that E2F1 regulates hepatic lipogenesis and bind directly with FASN promotor (Denechaud et al., 2016) and also the deficiency alleviate HCC in preclinical models (Gonzalez-Romero et al., 2021). In this context, p107 may modulate lipogenic programs through E2F-dependent networks. Further studies with E2F modulations are needed to define whether p107 exerts its metabolic effects through FASN. On the other hand it is important to note that these pocket proteins are classically described as regulators of the cell cycle, where they usually have redundant functions (Sage et al., 2003). In contrast, their roles in metabolism appear to be more distinct and no-redundant (Cuñarro et al 2019).

In summary, we identify p107 as a critical regulator of hepatic lipid metabolism. Its inhibition promotes a coordinated metabolic shift characterized by reduced DNL, decreased ER stress, and enhanced mitochondrial lipid oxidation, ultimately conferring protection against diet-induced steatosis in a FASN-dependent manner. These findings align with current knowledge (Loomba et al., 2021a) proposing that reversal of hepatic steatosis through metabolic interventions by improving hepatic lipid metabolism and prevent mitochondrial and peroxisomal dysfunction, inflammation, and progression to advanced liver disease. Within this context, p107 may provide a common nexus exploring its therapeutic potential, targeting p107 specifically in the liver could therefore offer a promising therapeutic strategy for metabolic liver disease.

## Graphical abstract

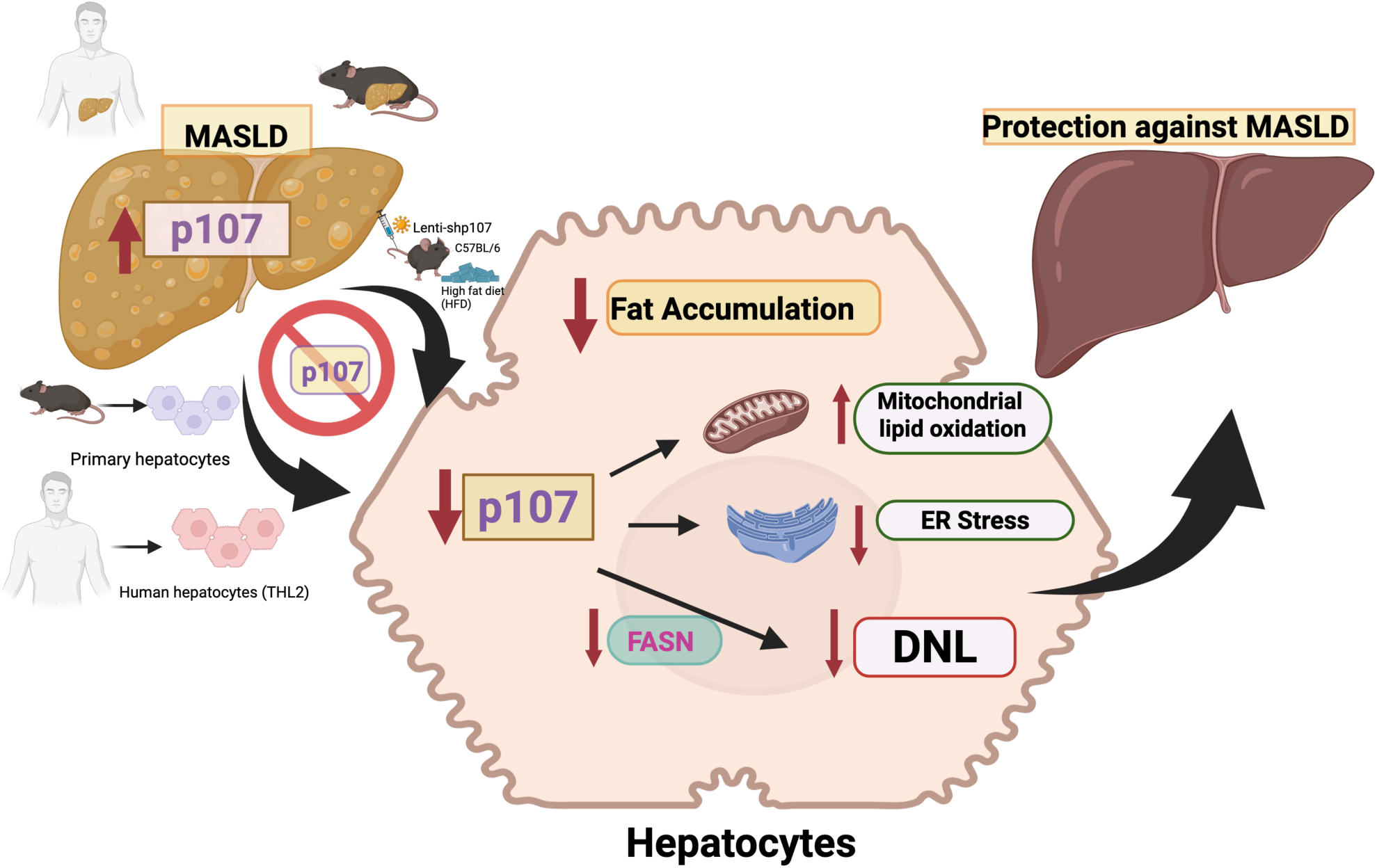

***Created in*** https://BioRender.com

## Acknowledgments

We thank Ana Senra for the help with histological analysis.

## DAS statement

Data available on request from the authors

## Author contributions

**Author contributions:** JC, MV-M designed and conducted experiments, analyzed results and draft the final figures and contributes to the manuscript preparation JC, MV-M TO-D, XB, LO, CQ-V, ATM, AC, EN AF-I performed the in vivo experiments and in vitro experiments. LF performed the proteomics and phosphoproteomics experiments, CR and JI. analyzed the proteomic experiments, JC, MV-M, RN, CD and ST contributes substantially to analysis and interpretation of the data. DG, MF, AV, AM writing, review, editing methodology and critically revising MS. MV-R, RN, GS, PA, LF, CD and ST drafting the work and interpreted and revising it critically for important intellectual content. ST formulated the hypothesis, secured the funding, coordinated the project and wrote the manuscript. ST are responsible for the integrity of the work as a whole.

---

**Supplementary Figure 1.**
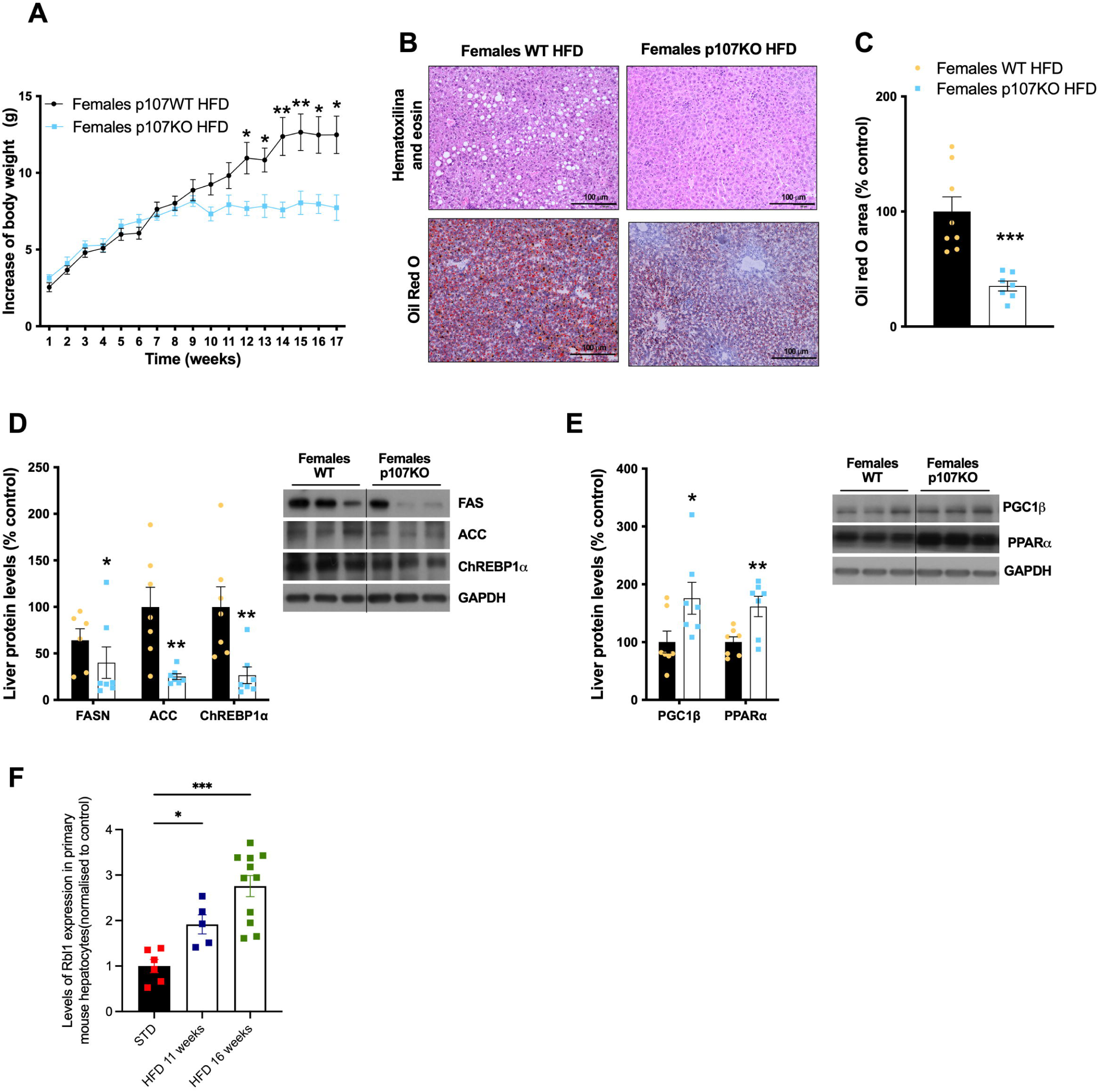
Female p107 KO mice are resistant to steatosis under very high-fat diet (HFD) and p107 levels in hepatocytes from mice with HFD. p107 KO and control female mice were fed a HFD for 16 weeks. (A) Body weight of control mice (n = 9) and p107 KO mice (n = 10). (B) Representative microphotographs of Oil Red O (upper panel) and H&E staining (lower panel) of liver sections. (C) Quantification of immunoblot analysis of *de novo lipogenesis* markers in liver (n = 7 per group) and a representative immunoblot. (D) Expression of *de novo lipogenesis* markers in liver (n = 8 per group). (E) Quantification of immunoblot analysis of lipolysis and mitochondrial activity markers in liver (n = 7 per group) and a representative immunoblot. GAPDH was used to normalize protein levels. (F) p107 expression in hepatocytes from liver’s mice with 11 or 16 weeks under HFD (n=6-11 per group). Data are expressed as mean ±SEM. *p < 0.05, **p < 0.01, ***p <0.001, using a Student’s *t test*.

**Supplementary Figure 2.**
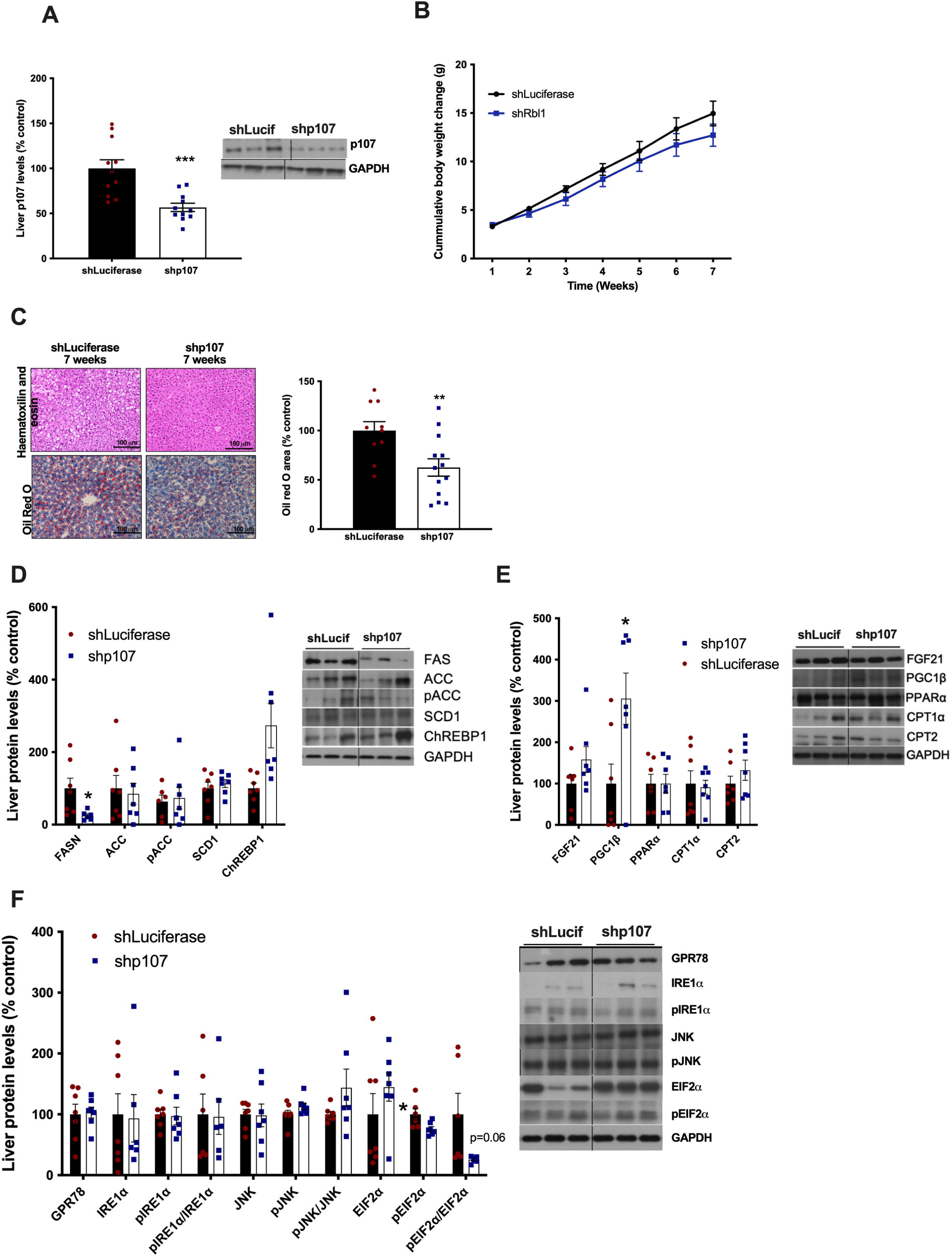
Liver-specific inhibition of p107 already produces resistance to hepatic fat accumulation at 7 weeks of HFD. Mice receiving the tail vein injection of lentivirus encoding shRNA p107 or luciferase were fed a HFD for 7 weeks. (A) Quantification of immunoblot analysis of expression of p107 in liver shRNA-Luciferase (n = 11) and in liver shRNA-p107 (n = 11) and a representative immunoblot. (B) Cumulative body weight change. (C) Representative microphotographs of H&E staining (upper panel) and Oil Red O (lower panel) of liver sections. Oil Red O staining was quantified using ImageJ (n = 10-13 per group). (D) Quantification of immunoblot analysis of *de novo lipogenesis* markers (n = 7 per group) and a representative immunoblot. (E) Quantification of immunoblot analysis of β -oxidation markers (n = 7 per group) and a representative immunoblot. (F) Quantification of immunoblot analysis of ER stress markers in liver (n = 7 per group) and a representative immunoblot. HPRT was used to normalize mRNA levels, and GAPDH was used to normalize protein levels. Data are expressed as mean ±SEM. *p < 0.05, **p < 0.01, ***p <0.001, using a Student’s *t test*.

