## Supplemental Table 1,2 and 3 for "Inhibition of p107 alleviates liver steatosis by reducing de novo fatty acid synthesis"

Supplementary Table 1: Anthropometric, biochemical and clinical characteristics of patients with normal liver (Non MASLD) and metabolic dysfunction-associated steatotic liver disease (MASLD) (n=24). Data are shown as mean ± standard deviation (SD) or as number of cases (%). ALT, alanine aminotransferase; AST, aspartate amino transferase; LDL, low-density lipoprotein.

| Feature | Non MASLD (n=12) | MASDLD (n=11) |
| --- | --- | --- |
| Age (years) | 48.0 ± 7.1 | 48.45 ± 11.7 |
| Body mass index (kg/m^2^) | 39.9 ± 6.4 | 41.9 ± 7.6 |
| Sex (%) |  |  |
| Men | 8 (66%) | 6 (54%) |
| Women | 4 (34%) | 5 (46%) |
| Body weight (kg) | 118.1 ± 26.3 | 117.0 ± 22.2 |
| Glucose (mg/dL) | 110.4 ± 54.7 | 133 ± 66.7 |
| LDL cholesterol (mg/dL) | 115.5 ± 34.7 | 108.8 ± 35.5 |
| Triglycerides (mg/dL) | 112.7 ± 55.2 | 181.7 ± 106.8 |
| Total cholesterol (mg/dL) | 191.3 ± 42.1 | 192.6 ± 50.0 |
| ALT (IU/L) | 28.8 ± 8.7 | 36.8 ± 17.4 |
| AST (IU/L) | 19.8 ± 9.6 | 23.8 ± 10.22 |
| NAS Score (%) |  |  |
| Grade 0 | 34 | 0 |
| Grade 1 | 58 | 0 |
| Grade 2 | 8 | 0 |
| Grade 3 | 0 | 36 |
| Grade 4 | 0 | 9 |
| Grade 5 | 0 | 46 |
| Grade 6 | 0 | 9 |
| Fibrosis (%) |  |  |
| Stage 0 | 92 | 8 |
| Stage 1 | 8 | 36 |
| Stage 2 | 0 | 56 |

Supplementary Table 2: List of primers used.

| Gene ID | Reference |
| --- | --- |
| hprt Mm | Taqman: Mm03024075_m1 |
| HPRT Hs | Taqman: Hs02800695_m1 |
| Rbl1 (p107) Mm | Taqman: Mm01250721_m1 |
| RBL1 (p107) Hs | Taqman: Hs00765700_m1 |
| Fasn (FAS) Mm | Taqman: Mm00662319_m1 |
| FASN (FAS) Hs | Taqman: Hs01005622_m1 |
| Acaca (ACC) Mm | Taqman: Mm01304257_m1 |
| Mlxipl (ChREBP) (Mm) | Taqman: Mm2342723_m1 |
| MLXIPL (Chrebp1) Hs | Taqman: Hs00975714_m1 |
| Srebf1 (Srebp1) Mm | Taqman: Mm00550338_m1 |
| Nr1h2 (LxR) Mm | Taqman: Mm00437265_g1 |

Supplementary Table 3: List of antibodies used.

| **Antibody** | **Reference** | **Host** | **Dilution** |
| --- | --- | --- | --- |
| **p107** | Abcam (ab244504) | Rabbit polyclonal | 1:1000 |
| **FAS** | Abcam (ab128870) | Rabbit monoclonal | 1:1000 |
| **ACC** | Abcam (ab45174) | Rabbit monoclonal | 1:1000 |
| **phospho-ACC (Ser79)** | Cell Signaling (3661) | Rabbit polyclonal | 1:1000 |
| **SCD1** | Cell Signaling (2794) | Rabbit monoclonal | 1:1000 |
| **SREBP-1c** | Abcam (ab3259) | Mouse monoclonal | 1:1000 |
| **ChREBP1a** | NOVUSBIO (NB400-135) | Rabbit polyclonal | 1:1000 |
| **FGF21** | Abcam (ab171941) | Rabbit monoclonal | 1:1000 |
| **PGC1β** | Santa Cruz Biotechnology (sc-373771) | Mouse monoclonal | 1:1000 |
| **PPAR α** | Santa Cruz Biotechnology (sc-398394) | Mouse monoclonal | 1:1000 |
| **CPT1A** | Abcam (ab128568) | Mouse monoclonal | 1:1000 |
| **CPT2** | Santa Cruz Biotechnology (sc-377294) | Mouse monoclonal | 1:1000 |
| **Citocromo C oxidasa** | Abcam (ab13575) | Mouse monoclonal | 1:1000 |
| **UCP-2** | Abcam (ab67241) | Mouse polyclonal | 1:1000 |
| **GPR78 (BiP)** | Cell Signaling (3183) | Rabbit polyclonal | 1:1000 |
| **ATF6**𝛂 | Santa Cruz Biotechnology (sc-166659) | Mouse monoclonal | 1:1000 |
| **IRE1**𝛂 | Abcam (ab37073) | Rabbit polyclonal | 1:1000 |
| **phospho-IRE1 (Ser 724)** | Novus Biologicals (NB100-2323) | Rabbit polyclonal | 1:1000 |
| **JNK 1/3** | Santa Cruz Biotechnology (sc-514539) | Mouse monoclonal | 1:1000 |
| **phospho-SAPK/JNK (Thr 183/Tyr 185)** | Cell Signaling (4668) | Rabbit monoclonal | 1:1000 |
| **XBP1** | Santa Cruz Biotechnology (sc-7160) | Rabbit polyclonal | 1:1000 |
| **eIF2α** | Santa Cruz Biotechnology (sc-11386) | Rabbit polyclonal | 1:1000 |
| **phospho-eIF2α (Ser 52)** | Santa Cruz Biotechnology (sc-101670) | Rabbit polyclonal | 1:1000 |
| **GADD 153 (CHOP)** | Santa Cruz Biotechnology (sc-793) | Rabbit polyclonal | 1:1000 |
| **GAPDH** | Merck -Millipore (G9545) | Mouse monoclonal | 1:5000 |
| **𝛂-tubulin** | Merck-Millipore (T5168) | Mouse monoclonal | 1:5000 |
| **Vinculin** | Santa Cruz Biotechnology (sc-73614) | Mouse monoclonal | 1:5000 |
| **HRP-conjugated anti-mouse**  **(secondary antibodies)** | Dako (P0260) | Rabbit polyclonal | 1:5000 |
| **HRP-conjugated anti-rabbit**  **(secondary antibodies)** | Dako (P0448) | Goat polyclonal | 1:5000 |
